# Tracking morphological development in stony corals

**DOI:** 10.1101/2025.11.20.689442

**Authors:** Garrett J. Fundakowski, Viviana Brambilla, Kyle J. A. Zawada, Cher F Y Chow, Emily Croasdale, Amelia J. F. Errington, Luisa Fontoura, Wilhelm J Marais, Rachael M. Woods, Pim Edelaar, Kevin Lala, Joshua S. Madin, Maria Dornelas

**Affiliations:** Centre for Biological Diversity, School of Biology, University of St Andrews, St Andrews, UK; MARE – Centro de Ciências do Mar e do Ambiente, Faculdade de Ciências, Universidade de Lisboa, Lisboa, Portugal; School of Natural Sciences, Macquarie University, Sydney, NSW, Australia; Reefscapers, Malé, Maldives; Australian Centre For Robotics (ACFR), University of Sydney, Sydney, NSW, Australia; Department of Climate Change, Energy, the Environment and Water, Sydney, NSW, Australia; Department of Molecular Biology and Biochemical Engineering, University Pablo de Olavide, Sevilla, Spain; Department of Biology, Lund University, Lund, Sweden; Hawai’i Institute of Marine Biology, University of Hawai’i at Mānoa, O’ahu, USA

**Keywords:** Morphological analysis, Scleractinia, 3D Photogrammetry, Traits, Growth form

## Abstract

The shape of reef-building corals largely determines how they interact with their environment and the ecosystem services they provide. However, morphology is not fixed. As corals grow and develop from singular polyps to mature adult colonies, they experience pronounced changes in their morphology. This developmental trait variation remains poorly quantified in corals, despite their importance as ecosystem engineers. To address this gap, we used photogrammetry to track the morphological development of wild coral colonies across six size-independent metrics designed to capture variation in volume compactness, surface complexity, and top-heaviness. Size-trait models and analyses of colony movement through morphological trait space revealed that small colonies shared compact morphologies and only later diverged along growth-form-specific developmental trajectories. Massive corals largely retained stable shapes as they grew, whereas non-massive colonies became less compact, more top-heavy, and in most cases, more complex. Despite these differences, there was still overlap between growth forms in their developmental trajectories through morphospace. These findings highlight that traditional categorical growth forms mask important developmental variation, particularly early in coral ontogeny. By applying a quantitative trait-based framework, this study improves understanding of coral morphological development and provides a foundation for linking individual growth trajectories to broader reef ecosystem dynamics.

## Introduction

Morphology plays a central role in mediating how organisms interact with their environment and contribute to ecosystem processes. However, morphology is dynamic – as individuals grow and develop, ontogenetic shifts in morphological traits can alter their performance and ecological processes. For example, size-related changes in leaf morphology can improve physiological performance by enhancing photosynthetic and water-use efficiency (Wang et al. 2025), while also reshaping ecological processes such as environmental filtering and community assembly (Spasojevic et al. 2014). Trait-based ecology has emerged as a powerful framework to study these links between form and function, but it has largely focused on interspecific differences, under the assumption that traits vary more between species than within them (McGill et al. 2006). Increasingly, studies have demonstrated that within-group trait variation can be substantial (Messier et al. 2010; Siefert et al. 2015) and that traditionally neglected drivers, such as temporal and developmental change, have important consequences for broader ecosystem dynamics (Bolnick et al. 2011; Westerband et al. 2021; Cope et al. 2022). Yet the developmental component of trait variation remains understudied, particularly in marine taxa where trait-based research lags behind that in terrestrial systems (Green et al. 2022).

Corals undergo considerable morphological changes during their lifespan and are a particularly important system for studying developmental trait variation because their morphology plays a critical role in reef functioning. Coral colonies begin their benthic life as a singular polyp, initially resembling a small hemisphere. As they replicate and the colony gets larger, their overall colony morphology differentiates and diverges across species. Some species develop into intricate, branching forms, while others form more dense, massive colonies. These ontogenetic changes in morphology influence both colony-level performance, such as vulnerability to hydrodynamic forces (Madin et al. 2014), and reef-level functioning, such as habitat provision for other organisms (Urbina-Barreto et al. 2021). Establishing links between colony size and shape is therefore important for improving our ability to understand and predict not only how individual colonies perform, but also how they collectively shape ecosystem-level processes on coral reefs.

Traditionally, Scleractinian corals are categorized into growth forms based on broad qualitative differences in adult colony morphology (Veron 2000). Growth form has been shown to be an important trait for delineating life-history strategies in corals (Darling et al. 2012), which predict their response to bleaching and fishing stressors (Darling et al. 2013). It has also been linked to differences in key demographic rates (mortality, Madin et al. 2014; fecundity, Álvarez-Noriega et al. 2016; growth, Dornelas et al. 2017). However, such a discrete categorization does not provide a complete description of a coral’s structure (Veron 2000) nor adequately captures the variation in colony shape both within and between species (Zawada et al. 2019a). In response to this shortcoming, and armed with recent advances in technology, many studies have begun quantifying the three-dimensional size and shape of coral colonies using photogrammetry, CT scanning, and laser scanning techniques (e.g., Bythell et al. 2001; Figueira et al. 2015; Gutiérrez-Heredia et al. 2015; Lavy et al. 2015; Reichert et al. 2016; House et al. 2018; Zawada et al. 2019a; Lange et al. 2020; Million et al. 2021; Urbina-Barreto et al. 2021; Guendulain-García et al. 2024).

Building on these advances, Zawada et al. (2019a) proposed a suite of six size-independent, morphological metrics that describe continuous three-dimensional shape space and position coral morphology along three functionally relevant axes of variation – volume compactness, surface complexity, and top-heaviness. Volume compactness, defined by sphericity and convexity, establishes a gradient from dense, boulder-like volumes to more intricate arborescent branching morphologies. Next, packing and fractal dimension describe surface complexity, spanning smooth to highly convoluted surfaces. Lastly, top-heaviness, characterized by the second moments of area and volume, describes the vertical distribution of colony surface area and volume relative to the substrate. These quantitative morphological traits have been linked to variation in organism performance, ecosystem function, and response to disturbance (Zawada et al. 2019b). While Zawada et al. (2019a) also established generic size relationships using these metrics, the corals they phenotyped were skeletons from museum collections and thus could not capture how individual colonies change through time. Thus, there remains a gap in understanding how these morphological traits develop through coral ontogeny, particularly in wild populations.

In this study, we address this gap by tracking morphological development in 133 wild coral colonies using the six quantitative metrics proposed by Zawada et al. (2019a). For each metric, we modelled size-trait relationships across all colonies and used these to examine how colonies move through multidimensional morphological trait space as they grow. We hypothesized that small corals from all growth forms would initially exhibit similar, compact morphologies before diverging along growth-form-specific developmental trajectories as they grow. Specifically, we expected volume compactness to be high in small colonies, with non-massive growth forms becoming less compact as they grow. We also anticipated surface complexity to be low in small colonies, with non-massive growth forms increasing in complexity (as these growth forms prioritize adding surface area over volume) and massive colonies maintaining their low values. Lastly, we predicted top-heaviness to be low in small colonies, increasing across non-massive growth forms as colonies grow, but most rapidly in tabular colonies. Overall, this study demonstrates the utility of these quantitative metrics for tracking coral morphology and improves our understanding of developmental trait variation in corals.

## Methods

### Study Site and Design

We conducted this observational study on ten reef sites around Jiigurru (also known as Lizard Island) at depths of 1-4 m in the northern Great Barrier Reef in Queensland, Australia (Fig. 1). To track morphological development in coral colonies *in situ*, we captured and quantified colony morphology of 133 coral colonies using photogrammetry over the course of four years and five fieldtrips (November 2018, November 2019, March 2021, April 2022, and November 2022). We selected colonies to capture a diverse array of species and morphological growth forms, ranging in size from small, young recruits to large, adult colonies. Exact locations of colonies were noted on printed orthomosaic maps to aid in colony relocation in subsequent years, though this was not always possible.

**Fig. 1.**
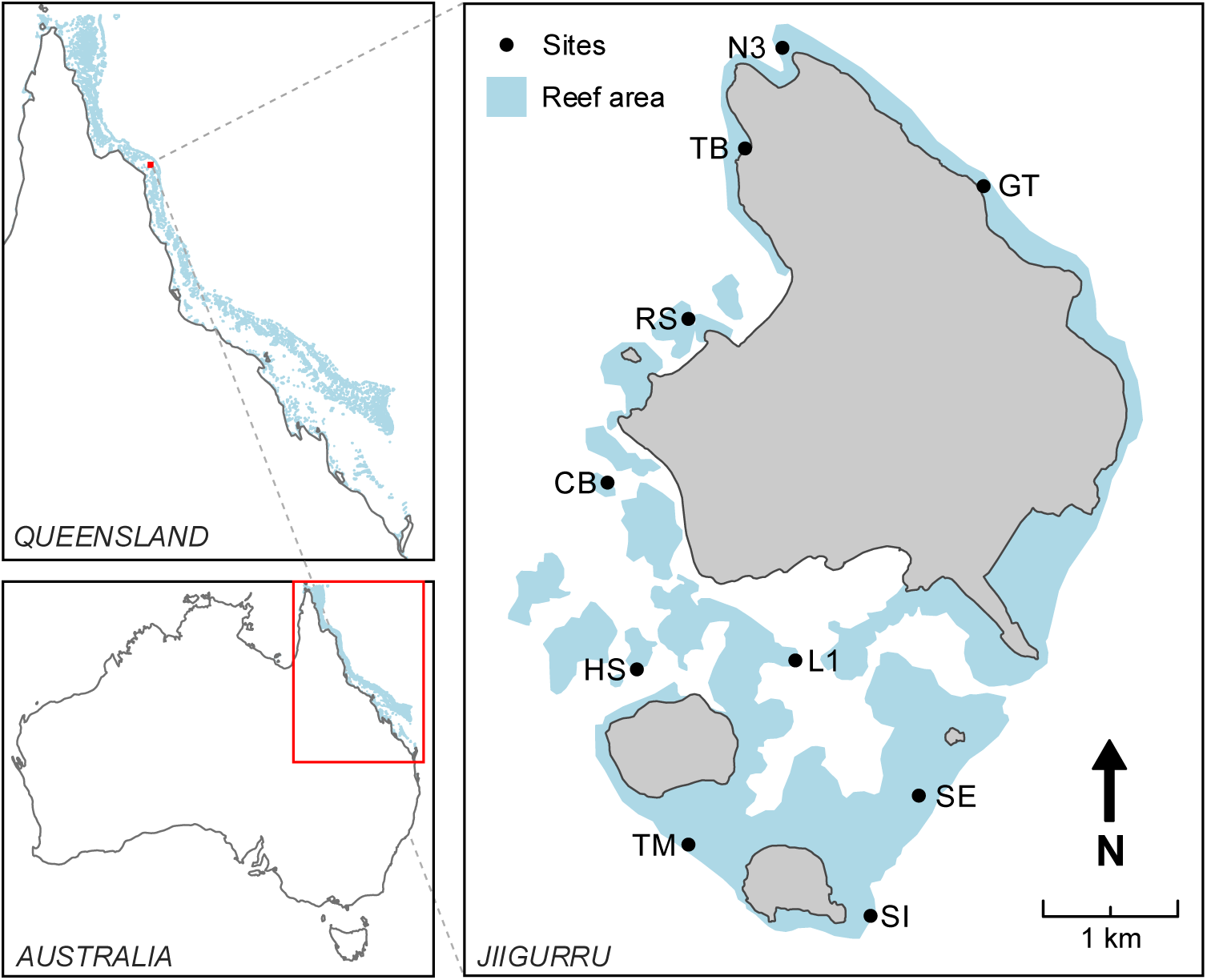
Map of study sites. The map shows the location of the 10 shallow water reef sites around Jiigurru, Australia. Moving counterclockwise from the north they are North Reef 3 (N3), Turtle Beach (TB), Resort (RS), Corner Beach (CB), Horseshoe (HS), Trimodal (TM), Lagoon 1 (L1), South Island (SI), Southeast (SE), and Gnarly Tree (GT). Spatial data for the Great Barrier Reef area and Jiigurru coastline are from Great Barrier Reef Marine Park Authority (GBRMPA) (2022)

### Image capture

We took photographs of selected colonies using handheld underwater cameras in continuous shooting/interval mode with autofocus enabled. We placed three gridded scale bars (2 cm x 2 cm or 10 cm x 10 cm) on the substrate surrounding the focal colonies to provide reference points for model scaling. Snorkelers photographed each reef scene (the area immediately surrounding the selected corals) while slowly swimming in a multi-tiered circular pattern about 0.5 m away from the focal colony. Photos were taken in an upward spiral pattern with views of the focal colony at approximately 0, 30, 60, and 80 degrees, maintaining roughly 60-80% overlap between images to ensure that the complex structure of the colonies was captured as completely as possible. Additional photos were taken of any complex sections that might require additional resolution to accurately build (e.g., dark undersides of tabular colonies or dense sections of branches in arborescent colonies). The number of images taken for each reef scene ranged from 73 to 1177 (average 298), with the number required for a good quality reconstruction increasing with size, complexity, and number of colonies in the reef scene being photographed.

### Model construction

We constructed three-dimensional (3D) models from the collected photosets using the proprietary photogrammetric software Agisoft Metashape Pro version 1.7.3 (Agisoft LLC 2021). The workflow used to generate the reef scene models for this study broadly follows the pipelines of similar studies that have used Metashape to construct 3D models from 2D images (e.g. - Lange et al. 2020; Million et al. 2021). However, meshes were built directly from depth maps (as in Miller et al. 2024) following recommendations from the Agisoft Metashape User Manual version 1.7 (Agisoft LLC 2021). We implemented this workflow with specified parameter settings (SI Table 1) using an automated Python script adapted from Million et al. (2021).

Following automated construction, we opened the reef scene models in Metashape for visual inspection and scaling. Any models with model construction issues (e.g. major deviations in shape, holes in the focal colony, or missing scale bars) were removed from the dataset. We then scaled the remaining models using four known lengths from two gridded scale bars, with the two lengths on the same scale bar being perpendicular to each other. We checked that scale error values were less than 1 mm and manually verified scaling accuracy by measuring the unused gridded scale bar. Finally, we exported the scaled reef scene models as a Wavefront (.obj) file with texture included, to allow for more precise cropping and isolation of the coral colony from the surrounding reef scene during post-processing.

### Method choice and validation

We chose to employ photogrammetry to measure colony morphology as it is currently the most accurate, cost effective, and non-invasive method available for doing so *in situ* (Figueira et al. 2015; Gutiérrez-Heredia et al. 2015). Multiple benchmarking studies have demonstrated that photogrammetric reconstructions yield surface area and volume estimates comparable to those obtained through CT, laser, and structured light scanning techniques (Figueira et al. 2015; Gutiérrez-Heredia et al. 2015; House et al. 2018; Guendulain-García et al. 2023). Further, a recent study from Miller et al. (2024) demonstrated the high precision and accuracy of a photogrammetric workflow using the same proprietary software and parameter settings. Across all reconstructed reef scene models in the current study, root mean square reprojection values ranged from 0.62 to 1.5 pixels, performing comparably to Miller et al. (2024) and indicating a high degree of 3D reconstruction accuracy in our models.

### Mesh post-processing

We then imported each scaled reef scene model into the free mesh-editing software Autodesk MeshMixer version 3.5 (Autodesk Inc. 2017) for post-processing to orient the reef scene, isolate the coral colonies, create a watertight mesh, and standardize mesh resolution.

First, we manually rotated the models to align the *z*-axis with the upwards facing direction of the reef scene *in situ* as some morphological metrics are orientation dependent. Models of the same reef scene from different years were oriented simultaneously to ensure consistency between years. Then we removed non-coral substrate from each reef scene model to isolate the focal individual(s). For scenes containing multiple focal colonies, we separated each colony into its own mesh file to allow for independent analysis.

At this point, the isolated colony meshes consisted of a continuous triangulated mesh with a hole where the colony was attached to the reef. We smoothed the boundary of this hole, evening out the jagged edges that were artifacts from the colony isolation process, to ensure all attachment boundaries were of similar smoothness regardless of mesh density. From here, we generated a new flattened attachment surface to create a watertight mesh, as this was necessary for calculating morphological metrics related to volume. To do this, we extended the mesh from the smoothed boundary in the direction of attachment to the reef (not always vertical). The extended mesh was then sliced using a plane, normal to the direction of extension, at the point where the plane intersected with the furthest extent of the smoothed attachment boundary. We filled the resulting cut with a flat plane to make the mesh watertight.

Finally, to ensure colony meshes were comparable, we standardized mesh resolution via a remeshing algorithm with a cell size of 2.4 mm, as this was the largest mean edge length across all colony meshes. The remeshing algorithm first creates an approximation of the input mesh using small cubes, or voxels, of the given cell size. The algorithm then reprojects the vertices from the voxel approximation back to the surface of the input mesh. In doing this, meshes retained their general shape with details smaller than the cell size being smoothed over, and resolution was standardized across meshes. Then we ran a final inspection of the mesh to fix non-manifold regions and fill small holes that may have been generated by the remeshing algorithm and exported the resulting watertight mesh in ASCII Polygon (.ply) file format. We automated the remeshing process and export of the final mesh with a custom Python script.

### Morphological metrics

To capture coral shape, we used the six morphological metrics described by Zawada et al. (2019a) – convexity, sphericity, packing, fractal dimension, and second moments of both area and volume – as they represent a quantitative, trait-based approach to describing coral morphology and have been shown to effectively partition quantitative morphological variation between qualitative growth forms. Convexity and sphericity define the morphological axis of volume compactness; packing and fractal dimension describe surface complexity; and second moments of both area and volume are top-heaviness metrics (Zawada et al. 2019a). Convexity, sphericity, packing, and fractal dimension are inherently size-independent metrics, while the second moments are not. Thus, to calculate the second moments of area and volume, meshes were first standardized by volume to remove the effect of colony size (Zawada et al. 2019a). In addition to these morphological traits, we extracted three fundamental size metrics from each colony mesh – planar area, surface area, and volume. All size and morphological metrics were extracted using the ‘habtools’ package (Schiettekatte et al. 2025) in R (R Core Team 2021). Specifically, to calculate fractal dimension we used the cube-counting method, as it is the recommended method for watertight 3D meshes (Schiettekatte et al. 2025).

### Colony Metadata

We identified each coral colony to species level sensu Veron (2000) and T. C. L. Bridge (personal communication). Then, we assigned growth forms for each species using the Coral Trait Database (CTD) (Madin et al. 2016b), which gives the ‘typical’ growth form of a species based on the morphological descriptions used in Veron (2000). Growth forms included were arborescent (referred to as open branching in the CTD), closed branching, corymbose, digitate, massive, submassive, and tabular (SI Table 2). For this study, we grouped the three colonies classified as submassives in with massives due to their similar morphologies.

The final dataset (Fundakowski et al. 2026) consisted of 340 watertight meshes from 133 unique colonies, with 104 colonies tracked across two or more years. The colonies spanned six growth forms, 36 species, and ranged in planar area from 1.03 cm^2^ to 2565.28 cm^2^ (see SI Table 3 for more details).

### Morphological development models

To investigate how morphological metrics change as corals grow, we fit six separate Bayesian hierarchical linear models with each metric as a response variable. To explore potential differences in the size-metric relationships across growth forms, we modelled colony shape as a function of the interaction between colony size and growth form for each morphological metric. For these analyses, planar area was used as the size covariate because it scales with 3D size traits (House et al. 2018) and is less biased than surface area or volume estimates to potential biases associated with generating watertight meshes.

Prior to model fitting, planar area was log_10_-transformed as the data spanned two orders of magnitude. Similarly, packing and both second moment metrics were also log_10_-transformed to account for the right-skewness in the data. Additionally, as sphericity and convexity are ratios bounded by zero and one, they were logit-transformed for analyses. With these transformations, response variables were approximately normal, and we modelled them with Gaussian distributions.

For each of the six models, we specified fixed effects to estimate growth-form-specific intercept 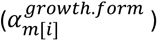 and slope 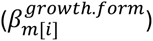 parameters with log_10_-transformed planar area (log_10_(𝑃𝐴_𝑖_)) as a continuous explanatory variable. We included Colony ID as a random intercept offset 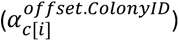 to account for any variation between colonies within growth form. We selected weakly regularizing priors for each parameter, broadly following McElreath (2020), and visually inspected them to ensure they did not constrain the posterior distributions (SI Fig. 1). The overall model parameterization and prior structure (Table 1) we implemented is as follows:

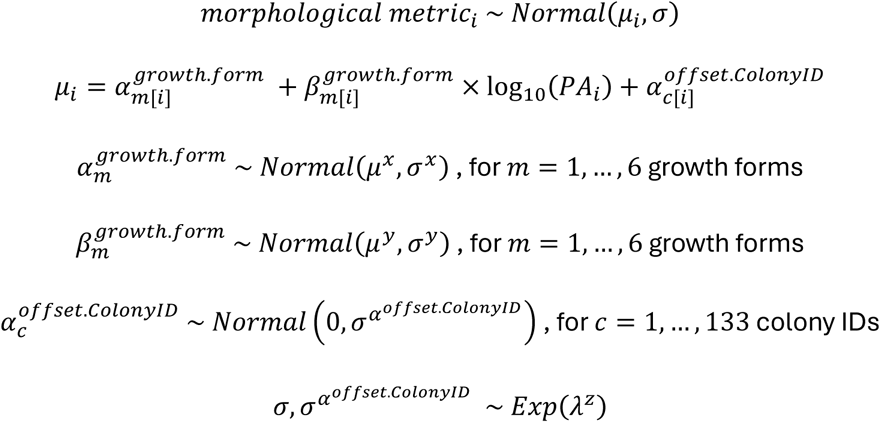

**Table 1.** Prior values used for each morphological development model.

| | $\alpha_m^{growth.form}$ | | $\beta_m^{growth.form}$ | | $\sigma, \sigma^{\alpha^{offset.ColonyID}}$ |
| --- | --- | --- | --- | --- | --- |
| *morphological metric<sub>i</sub>* | $\mu^x$ | $\sigma^x$ | $\mu^y$ | $\sigma^y$ | $\lambda^z$ |
| $\text{logit}(\text{convexity}_i)$ | 2 | 1 | 0 | 0.6 | 1 |
| $\text{logit}(\text{sphericity}_i)$ | 2 | 1 | 0 | 0.6 | 1 |
| $\log_{10}(\text{packing}_i)$ | -0.2 | 0.4 | 0 | 0.2 | 3 |
| $\text{fractal dimension}_i$ | 2.2 | 0.2 | 0 | 0.2 | 3 |
| $\log_{10}(\text{second moment of area}_i)$ | 0 | 0.5 | 0 | 0.2 | 3 |
| $\log_{10}(\text{second moment of volume}_i)$ | -0.5 | 0.5 | 0 | 0.2 | 3 |

### Growth models

We fit a second set of Bayesian hierarchical models to quantify changes in three fundamental size metrics – planar area, surface area, and volume – through time. For each size metric, we used Gaussian distributions to model colony size at one timepoint as a function of size at the previous timepoint and its interaction with growth form. The model structure is broadly similar to the morphological development models and included growth-form-specific intercept 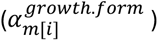 and slope 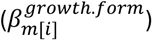 parameters, as well as a random intercept offset 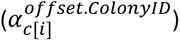 to account for ColonyID. As before, we assigned weakly regularizing priors for each parameter, with slopes centered around 1 and intercepts around 0 to represent no net change in size, and visually inspected them to ensure they did not constrain the posterior distributions (SI Fig. 2). The overall model parameterization and prior structure we implemented is as follows:

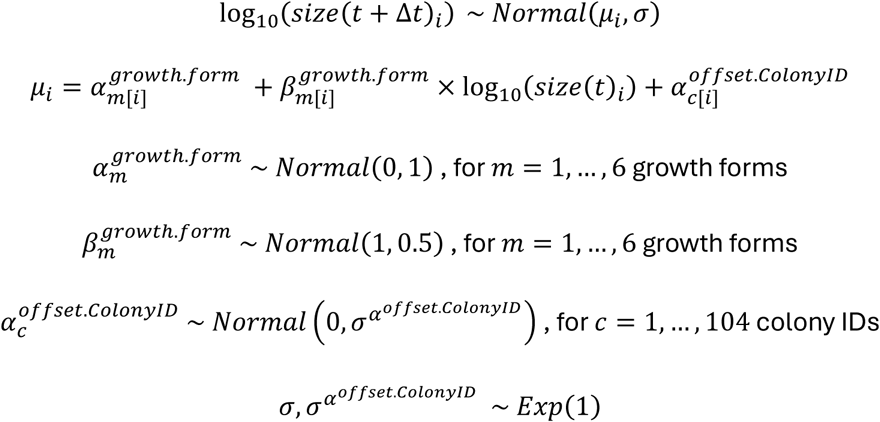

### Model fit and validation

We fit all models in R (R Core Team 2021) using stan statistical programming language (Stan Development Team 2024) via the ‘brms’ package (Bürkner 2017). We ran four MCMC chains of 20000 iterations per chain, with a warm-up of 5000 iterations and a thinning rate of 10 iterations, for each model. This resulted in 6000 draws in the posterior distributions. To assess model convergence, we confirmed *R̂* values were sufficiently close to 1.

To check model accuracy in reproducing the empirical data, we performed a set of posterior predictive checks for each model (Gelman et al. 1996; Gabry et al. 2019; Gelman et al. 2020). First, we produced a set of 1000 simulated datasets using the ‘posterior_predict’ function from the ‘brms’ package (Bürkner 2017). Then we calculated the mean and standard deviation of the response variable from these simulated datasets and compared them to the summary statistics from the raw data across varying grouping levels to assess model fit. We did these checks at both the growth form and site level for all morphological development models (SI Fig. 3 – SI Fig. 8) and growth models (SI Fig. 9 – SI Fig.11).

When reporting model results, we borrow the language of evidence as proposed by Muff et al. (2022).

### Developmental trajectories in morphological trait space

We used Principal Component Analysis (PCA) to construct a morphological trait space (morphospace) from the six morphological metrics derived from colony meshes. Prior to ordination, each metric was standardized to have a mean of zero and unit variance to remove the bias from varying metric scales. We conducted PCA significance tests using the ‘PCAtest’ package (Camargo 2022) to determine the significant principal components (PCs) (*p* < 0.05) and identify metrics with significant loadings on each PC.

To examine how colonies developed within this morphospace, we extracted fitted values from each of the six morphological development models and projected them onto the PCA space. To get colony-level trajectories, we drew 6000 samples from the expected values of the posterior predictive distribution across the range of observed planar areas, including ColonyID as a random effect for all 104 colonies tracked for at least two years. From these draws, we calculated colony-level displacement vectors as the difference between the predicted positions at the minimum and maximum sizes (standardized across growth forms), and we summarized the distribution of vector magnitudes and directions by growth form. Finally, to compare how morphologies occupied morphospace at different size classes, we randomly sampled 100 posterior predictive draws per colony at 15 standardized planar areas, calculated from a set of colony diameters assuming circular geometry, and plotted the density of predicted positions by growth form.

## Results

### Morphological development models

All growth forms except for massives exhibited strong evidence (99% credible interval below zero; Muff et al. 2022) for negative size-compactness relationships across both convexity and sphericity models (Fig. 2, SI Table 4). Massives had a slope estimate distribution that overlapped with zero suggesting they maintained relatively constant volume compactness across the observed size range. In the convexity model, tabular, digitate, and arborescent colonies had steeper slope estimates than the closed branching and corymbose colonies. For the sphericity model, the slope estimates for these non-massive growth forms were broadly similar, with digitate colonies having a slightly more negative slope estimate.

**Fig. 2.**
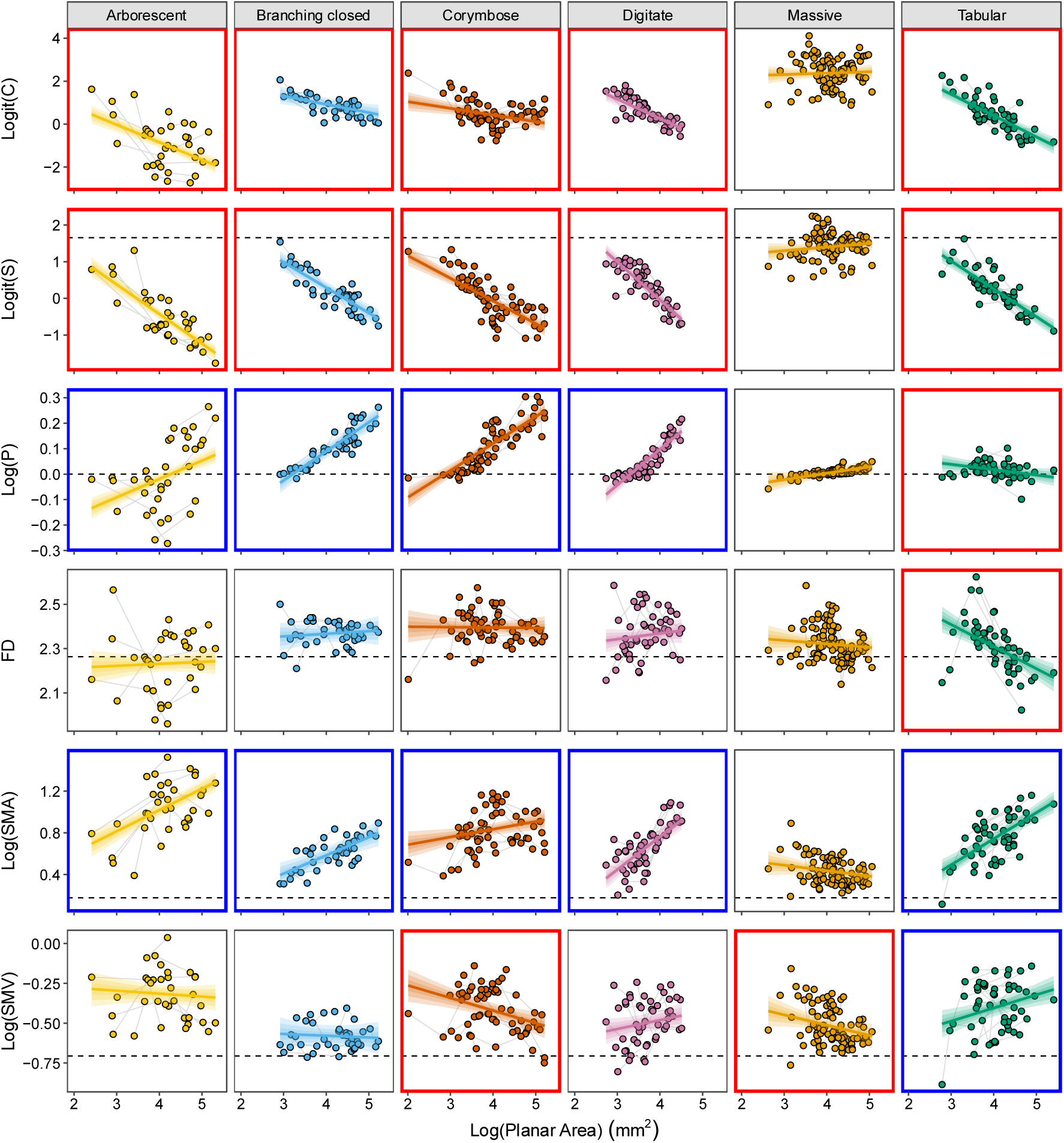
Scatterplots for all morphological development models. Each row corresponds to a different morphological development model (C = Convexity, S = Sphericity, P = Packing, FD = Fractal dimension, SMA = Second moment of area, SMV = Second moment of volume) faceted by growth form. Here, points represent individual meshes (n = 340) colored by growth form, with grey lines connecting the meshes from the same Colony ID through time. Bold colored lines represent linear regression lines (with 50%, 80%, and 95% credible intervals) overlaid for each growth form. Black horizontal dashed lines represent corresponding metric values for a hemisphere. (Note: a perfect hemisphere has a convexity value of 1, which on the logit scale would be positive infinity and therefore is not displayed.) Individual plot borders are colored based on whether the 95% credible interval for model slope estimates overlapped with zero for each growth form (red = negative slopes, grey = slopes overlapping zero, blue = positive slopes)

Size-complexity relationships displayed distinct patterns across growth forms depending on the surface complexity metric considered. In the packing model, arborescent, closed branching, corymbose, and digitate colonies showed strong evidence for positive size-packing relationships while table corals demonstrated strong evidence for a slight negative relationship across the observed size range. Massives showed weak evidence (90% credible interval above zero; Muff et al. 2022) for a positive size-packing relationship. On the other hand, for the fractal dimension model, table corals were the only growth form to show strong evidence for a size-fractal dimension relationship, which was negative. All other growth forms did not experience a change in their fractal dimension with increasing size, as their credible intervals all overlapped zero.

With regard to the top-heaviness models, both second moment of area (SMA) and second moment of volume (SMV) exhibited non-zero relationships for some growth forms. In the SMA model, arborescent, closed branching, digitate, and tabular colonies all showed strong evidence for positive slopes, while corymbose colonies exhibited moderate evidence (95% credible interval above zero; Muff et al. 2022). Among these, digitate and tabular colonies had the steepest positive slopes. Massives, however, maintained a constant SMA as size increased, with the credible interval of the slope estimate overlapping zero. The relationships were quite different for the SMV model. Here, table corals exhibited strong evidence and digitate colonies exhibited weak evidence for increasing SMV with increasing size. On the other hand, corymbose colonies displayed strong evidence and massives displayed moderate evidence for a negative relationship between size and second moment of volume. Meanwhile, arborescent and closed branching corals had slope estimates overlapping zero, indicating no detectable change in SMV with size.

### Growth models

For the most part, all growth forms across all growth models exhibited moderate (95% credible interval) to strong (99% credible interval) evidence of positive slopes less than one and positive intercepts (Fig. 3 Scatterplots for all growth models.Fig. 3, SI Table 5). The one exception was the digitate growth form for the planar area growth model which showed only weak evidence (90% credible interval) for a positive slope estimate less than one. Across all growth models, massive colonies had the slowest and arborescents had the fastest growth rates for the observed size range, with the other growth forms changing rank order with increasing size. Together, these parameter estimates define regressions that cross the unity line, a line on the log-log scale with a slope of 1 and intercept of 0, which corresponds to no change in size between the two timepoints. Points above this line represent individuals with positive net growth, and those below it, negative net growth (i.e. they shrunk). The medians from the posterior distributions of these intersection points showed a consistent ordering across all growth models. Massive colonies cross the no net growth on average line at the smallest sizes, followed closely by closed branching and table corals (and, for volume, digitate colonies too), while arborescent and corymbose growth forms crossed the 1:1 line at the largest sizes.

**Fig. 3.**
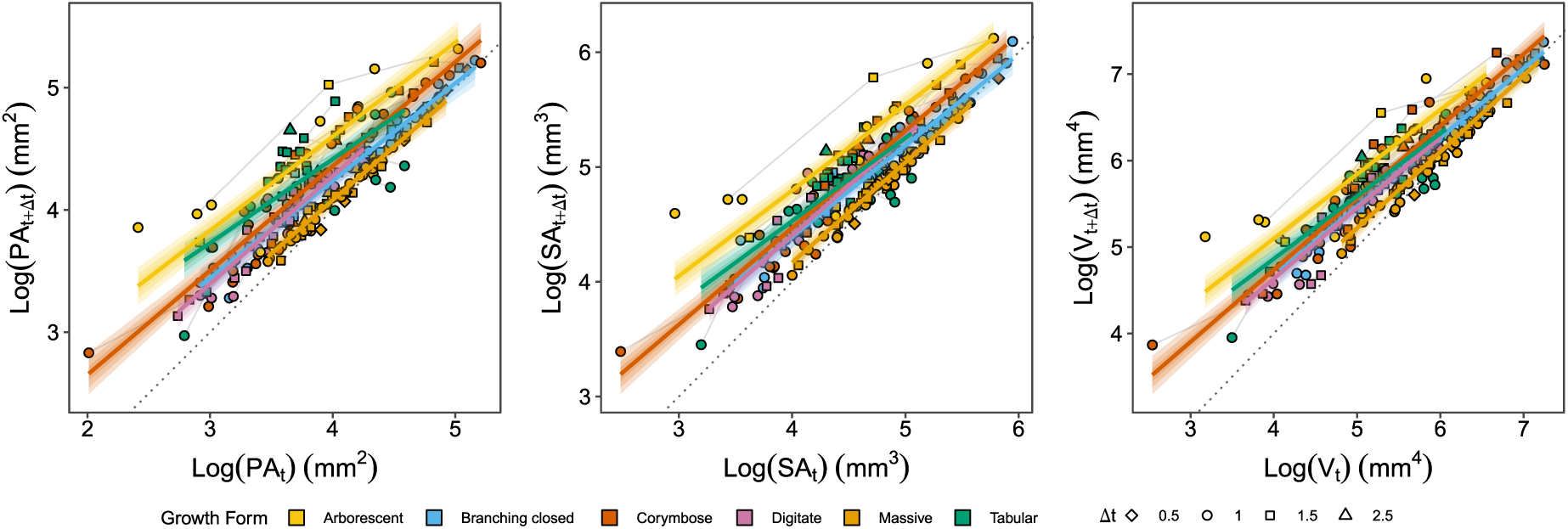
Scatterplots for all growth models. Each plot corresponds to a different size metric used (PA = Planar Area, SA = Surface Area, V = Volume). Here, points represent paired size observations (n = 207) for an individual colony, linking a mesh at time 𝑡 to the corresponding colony mesh at time 𝑡 + Δ𝑡. Points are colored by growth form with their shape corresponding to Δ𝑡, and grey lines connect successive paired observations from the same Colony ID through time. Bold colored lines represent linear regression lines (with 50%, 80%, and 95% credible intervals) overlaid for each growth form. Black diagonal dashed lines mark the line of no change

**Fig. 4.**
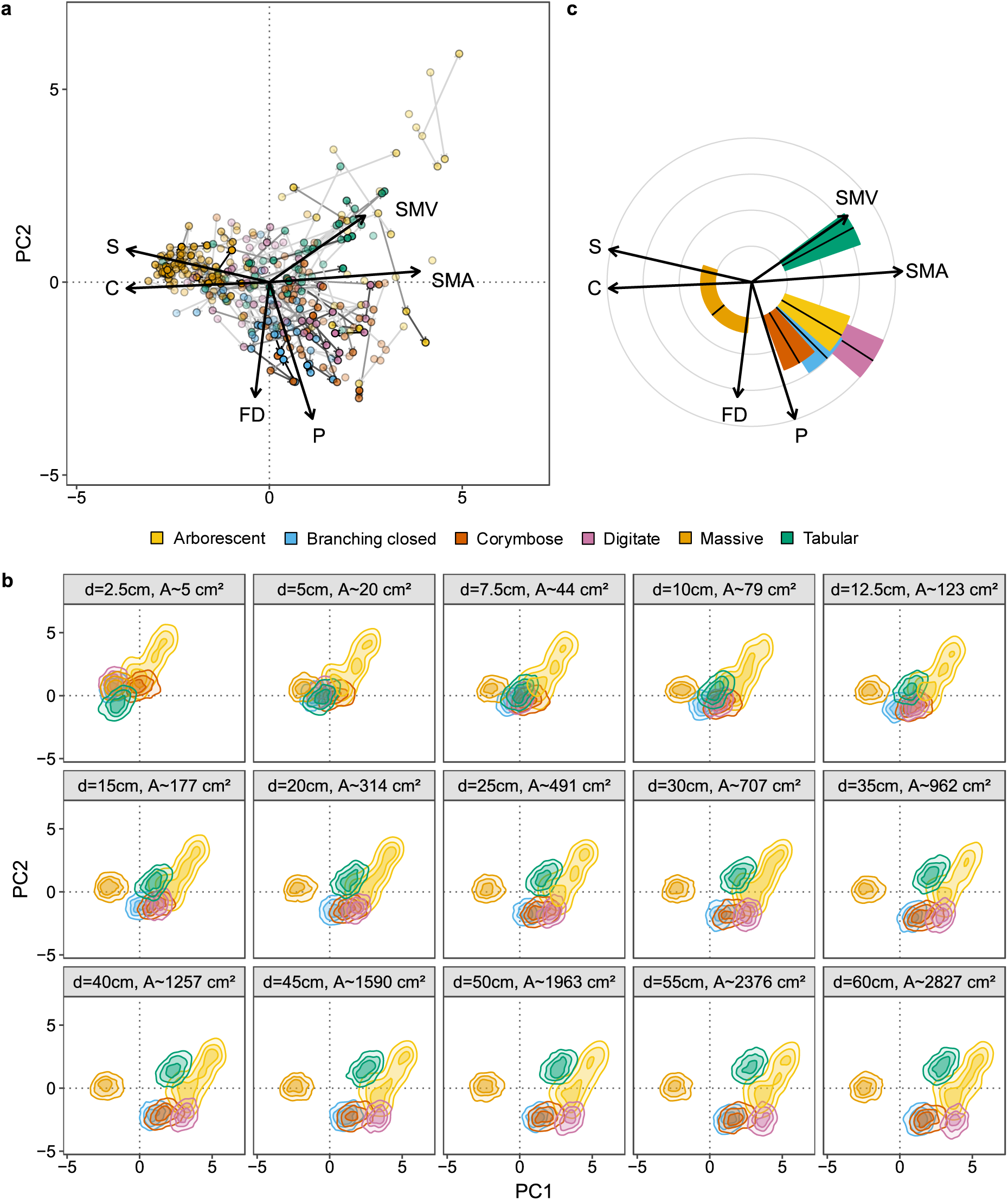
Morphological development in continuous trait space. The multidimensional space was built using the six morphological metrics (C = convexity, S = sphericity, P = packing, FD = fractal dimension, SMA = second moment of area, SMV = second moment of volume) extracted from the colony mesh models. The space is projected in two dimensions using the first two principal components (PC), which account for 78.3% of the total variance. (a) Points depict individual meshes (n = 340) colored by growth form. Meshes from the same colony are connected by grey arrows, with the direction and the increasing opacity of both points and arrows indicating progression over time. Black arrows represent the loading and direction of each metric with arrow lengths showing how much each trait contributes to the PCs (scaled by a factor of 4 for visualization). (b) Normalized density contour plots representing the portion of the total morphospace occupied by each growth form at various colony diameters and corresponding circular planar areas. (c) Circular plot depicting the direction and magnitude of the mean trajectory (solid black line) for each growth form. Colored bars represent each growth form with their heights showing the relative magnitude of the mean trajectory and their widths depicting the 95% quantile range for the direction of individual colony trajectories. Black arrows represent each metric as it loads into the PCA

### Developmental trajectories in morphological trait space

When defining the morphological trait space, the first two principal component (PC) axes were significant (*p* < 0.001) and together captured 78.3% of the observed variation in shape (Fig.4a). The first PC explained 52.1% of the variation across the morphospace and was comprised of the four volume compactness and top-heaviness metrics. The second PC captured 26.2% of the variation and was defined by the two surface complexity metrics.

Model predictions showed that small colonies across all growth forms were concentrated in the region of the morphospace associated with higher volume compactness (Fig.4b). With increasing planar area, the areas occupied by each growth form progressively diverged. At approximately 45 cm^2^ (equivalent to a circle with diameter 7.5 cm), a division appeared between massive and non-massive growth forms, with the latter shifting positively along PC1, in the direction of decreasing compactness and increasing top-heaviness. As size increased further, additional differentiation occurred among non-massives, particularly along PC2, reflecting increasing variation in surface complexity. Here, at approximately 950 cm^2^ (35 cm diameter circle) arborescent and tabular colonies largely occupied regions of the morphospace distinct from other growth forms. In contrast, digitate colonies did not become reliably distinguishable until the largest size class considered (approximately 2800 cm^2^, 60 cm diameter circle), while corymbose and closed branching colonies sustained a high degree of overlap.

Looking at how coral colonies moved through the morphological trait space, there was a strong pattern of growth form differences in their developmental trajectories (Fig.4c, SI Table 6). Here, massive colonies clearly stood out from the other growth forms with the smallest magnitude mean trajectory vector, suggesting relatively consistent morphology with increasing planar area. Massives were also the only group to shift on average toward increased volume compactness and decreased top-heaviness (moving negatively along PC1). However, it is also worth noting they exhibited the greatest colony-level variation in trajectory direction compared to other growth forms. Table corals also had a distinctive trajectory, as the only growth form moving toward reduced surface complexity (moving positively along PC2). The rest of the growth forms, on the other hand, generally followed trajectories of decreasing compactness (moving positively along PC1) and increasing surface complexity (moving negatively along PC2), though subtle differences in magnitude and direction were evident. For instance, corymbose, closed branching, and arborescents exhibited similar magnitudes of change, but differed slightly in direction, with corymbose colonies shifting more strongly toward greater surface complexity (negatively along PC2). Meanwhile, digitate colonies paralleled arborescents in trajectory direction but had a much greater trajectory magnitude across the standardized range of planar area values.

## Discussion

Using a quantitative framework to describe complex three-dimensional (3D) structure, this study tracked the morphological development of coral colony growth *in situ*. We used six metrics that describe colony geometry and morphology and explicitly showed the divergence and similarities between nominal growth forms throughout development. As predicted, corals of all growth forms initially exhibited similar compact morphologies, beginning as small semi-hemispherical colonies characterized by high volume compactness, low surface complexity, and low top-heaviness. As they grew, corals belonging to different growth forms took different developmental trajectories through morphological trait space, with non-massive growth forms showing decreases in both sphericity and convexity, as well as increases in second moment of area with size. Although divergence among growth forms gradually became more apparent with increasing size, there was still considerable overlap between some growth forms even at the largest sizes considered. This overlap suggests that traditional qualitative growth forms for corals may only be useful categorizations for large mature colonies, as at smaller sizes there is too much overlap in morphological space to reliably distinguish among growth forms. Furthermore, these findings highlight the complexity of morphological development and underscore the importance of quantitative approaches for capturing the full range of morphological variation.

One of the clearest departures from predicted size-trait relationships occurred in table corals, particularly in metrics describing surface complexity. While most non-massive growth forms followed the expected positive size-packing patterns, tabular colonies instead exhibited a slight negative size-packing relationship. Despite this, their packing values remained close to 1 (log-packing values centered around 0) across the observed range of sizes. A packing value of 1 indicates that the surface area of the object is equal to that of its convex hull, a feature obviously achieved by an entirely convex object. Although table corals are not fully convex, such packing values can also result when an object has a balanced amount of inverted and everted surfaces (Zawada et al. 2019a), and the model results suggest that table corals experience a subtle reduction in this ratio as they grow. This observed negative size-packing relationship in table corals was reinforced by an also unexpected negative relationship between size and fractal dimension. However, previous research has revealed that as table corals get larger, the shelter volume (e.g. unoccupied volume under the table coral) they provide increases exponentially (Urbina-Barreto et al. 2021), suggesting that the corals themselves occupy a progressively smaller fraction of the bounding box. It would make sense then, that for the cube-counting algorithm employed to estimate fractal dimension, that with increasing colony size, increasingly fewer cubes will intersect the coral surface, leading to the observed negative relationship. Thus, although initially unexpected, both packing and fractal dimension results converge on a consistent result – that table corals lose surface complexity as they grow.

By contrast, non-tabular morphologies maintained a relatively constant fractal dimension with increasing size, rather than the predicted positive relationship. Notably, across all morphological development models, fractal dimension produced some of the highest residuals suggesting that some variation remained unexplained. This may be partly attributable to methodological constraints. To account for different mesh resolutions resulting from variable imaging conditions, all colonies were remeshed to a standardized resolution prior to metric extraction. While necessary to ensure comparability between meshes, this step unavoidably introduced bias by smoothing out fine-scale surface texture, likely reducing fractal dimension estimates and potentially obscuring biological patterns. Though other 3D modelling technologies (i.e. laser-scanning or CT scanning) can achieve higher resolution and capture finer-scale detail than 3D photogrammetry (Figueira et al. 2015; Guendulain-García et al. 2023), these systems are prohibitively expensive. Thus, at the moment, underwater photogrammetry is the most accurate and feasible non-invasive method available for obtaining *in situ* morphological trait measurements (Figueira et al. 2015; Gutiérrez-Heredia et al. 2015).

Unexpected results were also observed in the relationship between size and second moment of volume (SMV). Instead of the predicted increase in SMV with size in all non-massive corals, several growth forms showed either no change or a decrease in SMV with increasing size. One possible explanation lies in the manual isolation step required to extract colonies from the surrounding reef scene. By trimming the colony at the living tissue edge and extending boundaries toward the attachment point, portions of the underlying substrate that the colony grew over (rather than created itself) may have been inadvertently included. This effect likely varies by growth form, with larger, broad-based colonies gaining disproportionately more volume during processing than smaller colonies with narrow attachment points. Such methodological bias could obscure true size-SMV relationships and likely contributed to the high residual variation observed across growth forms as well. Approaches that align dense point clouds between years before colony isolation could help standardize trimming (as in Reichert et al. 2016) and may reduce this source of variation, albeit at the cost of increased processing time and additional software requirements.

Importantly, the developmental trajectories observed in morphological trait space must also be interpreted in the context of net colony growth. In line with the findings from Dornelas et al. (2017), our growth models revealed all growth forms exhibit allometric growth, with slopes consistently less than one in log-log space indicating that colonies increase in size up to a growth-form-specific point. By explicitly modelling these changes in planar area, surface area, and volume through time, we showed that growth dynamics vary among growth forms and partially – but not entirely – explain differences in growth-form-level trajectories through morphospace. Massive colonies, which exhibited the slowest growth rates across all size metrics, also displayed the smallest magnitude trajectories through morphological space, consistent with their relatively conserved morphology through ontogeny. In contrast, arborescent colonies grew the fastest, yet did not exhibit the largest magnitude movement through morphospace, indicating that rapid growth does not necessarily translate into proportionally large movements through morphospace. Conversely, digitate colonies displayed the largest magnitude trajectories in morphospace despite intermediate growth rates, suggesting that relatively modest increases in size can be accompanied by substantial changes in colony morphology. Together, these patterns demonstrate that developmental trajectories through morphospace are less about how fast corals grow, and more about how that growth is allocated across 3D space. However, growth-form-level differences in skeletal density or calcification rates (Hughes 1987) - traits not explicitly measured here – might possibly explain these different trajectories and represent important avenues for future work.

The finding that small colonies are morphologically similar across growth forms raises the question of at which size do corals begin to diverge into distinct growth-form-specific morphologies. Instead of a single threshold size, our results reveal that morphological divergence was gradual, occurring along different axes of shape variation at different sizes. This suggests that growth form differentiation is a continuous developmental process rather than a discrete transition. However, it is important to note, that threshold size will depend on the context – the traits considered, the resolution of morphological measurement, and the ecological function of interest. For example, distinctions relevant to mechanical stability (Madin and Connolly 2006) may emerge at smaller sizes than those related to fish shelter provision (Urbina-Barreto et al. 2021). As such, rather than identifying a universal size threshold, our results emphasize the need to consider morphology as a continuum when linking coral size, shape, and ecological function.

The trajectories of morphological development observed in this study have important ecological and evolutionary implications. As coral colonies grow and change shape, their impact on the environment and local reef community will change too. For instance, bigger corals will absorb more light and cast larger shadows (Stambler and Dubinsky 2005), thus altering the light availability for sessile benthic organisms in their vicinity. Additionally, as table corals grow and they become larger and more top-heavy, they can offer greater shelter volume (Urbina-Barreto et al. 2021), which is critical for large-bodied reef fish (Kerry and Bellwood 2011,2014). On the other hand, as corals grow, their changing shape also alters how they experience their environment, and in turn the selection they are exposed to. For example, larger table corals are more top-heavy than smaller ones, making them more prone to dislodgement and damage from wave energy (Madin and Connolly 2006; Zawada et al. 2019b). This selective impact of hydrodynamic disturbances on the size structure of tabular species has potential downstream implications for population recovery, as the most mechanically vulnerable tabular colonies are the most fecund (Álvarez-Noriega et al. 2016). By tracking the morphological development of corals using quantitative metrics, this study deepens our understanding of how coral growth might shape ecological interactions in reef ecosystems.

While the results of this study capture some growth-form-specific size-metric relationships and trajectories through continuous morphological space, they also emphasize the large amount of overlap between growth forms in morphological trait space, echoing the findings of Zawada et al. (2019a). However, the current results reveal even more extensive overlap in morphological traits across growth forms, specifically highlighting the developmental variation of coral morphology both within an individual colony and between individuals of a given growth form. This reinforces the idea that these traditional growth forms lack the clear boundaries implied by their categorical nature (Zawada et al. 2019a), especially at smaller colony sizes. The intra-individual and within-growth-form developmental variation of coral morphology observed here once again underscores the need to transition from qualitative to quantitative trait-based approaches to better understand fundamental biological and ecological processes on reefs (Madin et al. 2016a).

By elucidating these size-metric relationships and individual morphological development, this study lays the groundwork for future studies to more explicitly test reciprocal interactions between coral morphology and reef ecosystem processes. This quantitative framework can be extended to directly evaluate how morphological changes impact the surrounding environment, and how environmental conditions in turn shape coral morphology, working to establish causal links between the two processes (Jones et al. 1997; Odling-Smee et al. 2003). Integrating this morphological data with environmental variables like light availability, temperature, and water movement for example, offers a promising avenue for understanding the feedback loop between coral structure and the reef environment.

## Data availability

Python code to generate the 3D meshes in Agisoft Metashape as well as the final 3D mesh dataset can be accessed at: https://doi.org/10.17630/b0086b3f-fec9-4765-9a0f-fe9021aa79e3 (Fundakowski et al. 2026). Data and code required to reproduce the analyses in this paper can be accessed at: https://doi.org/10.17630/b562de9c-f339-4694-9e07-44c04f870c5d (Fundakowski 2026).

## Statements and Declarations

The authors of this paper declare that there is no conflict of interest.

## Author contributions

Conceptualization: GJF, VB, KJAZ, JSM, MD. Funding Acquisition: VB, KL, JSM, MD. Resources: JSM, MD. Supervision: PE, KL, JSM, MD. Investigation: GJF, VB, KJAZ, CFYC, LF, WJM, RMW, MD.

Methodology: GJF. Data Curation: GJF, EC, AE. Formal analysis: GJF. Writing – original draft: GJF. Writing – review and editing: all authors.

## Acknowledgements

We thank the Australian Museum’s Lizard Island Research Station staff for their support in facilitating this research. This study was conducted under GBRMPA research permits G15/38127.1 and G22/46527.1. We would also like to thank Mariana Álvarez-Noriega, who assisted with data collection in March 2021, as well as Devynn Wulstein, Nina Schiettekatte, and Mollie Asbury, who were members of the April and November 2022 field teams.

Funding was provided by the John Templeton Foundation (grant #60501 ‘Putting the Extended Evolutionary Synthesis to the Test’) (KL, MD, JSM), an Ian Potter Doctoral Fellowship at Lizard Island Research Station (VB), a MASTS small grant (VB), a Leverhulme fellowship (MD), University of Hawai’i start-up funding (JSM), the Warman Foundation (MD, JSM), a National Science Foundation–Natural Environment Research Council Biological Oceanography Grant (1948946) (JSM, MD), and the European Union (CoralINT, GA 101044975) (MD, VB). Views and opinions expressed are however those of the author(s) only and do not necessarily reflect those of the European Union or the European Research Council. Neither the European Union nor the granting authority can be held responsible for them.

