## Supplemental Information for "Tracking morphological development in stony corals"

### Supplementary Information

**SI Table 1** Parameter settings used during 3D model construction in Agisoft Metashape

| **Process** | **Parameters** |
|---|---|
| Image Quality | Image quality threshold: 0.5 |
| Align Photos | Accuracy: High<br>Generic preselection: Yes<br>Reference preselection: Source<br>Key point limit: 40,000<br>Tie point limit: 1,000<br>Guided image matching: No<br>Adaptive camera model fitting: Yes |
| Gradual Selection | Reconstruction uncertainty threshold: 30<br>Projection accuracy threshold: 10<br>Reprojection error threshold: 0.7<br>Camera optimization: f, cx, cy, k1–k3, p1–p2 |
| Build depth maps | Quality: High<br>Filtering mode: Mild |
| Build mesh | Surface type: Arbitrary<br>Source data: Depth maps<br>Interpolation: Enabled<br>Face count: High<br>Calculate vertex colors: Yes<br>Reuse depth maps: Yes |
| Build texture | Mapping mode: Generic<br>Blending mode: Mosaic<br>Texture size: 16,000<br>Enable hole filling: Yes<br>Enable ghosting filter: Yes |

**SI Table 2** Growth form examples and descriptions

| | | |
|---|---|---|
| **Arborescent** | 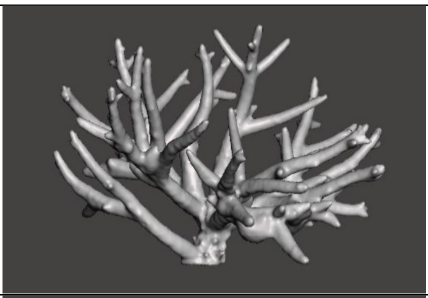 | 'Tree-like' growth form with lots of open space between branches |
| **Closed Branching** | 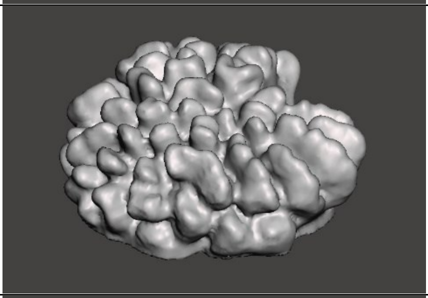 | Growth form with branches that form in clusters or tufts |
| **Corymbose** | 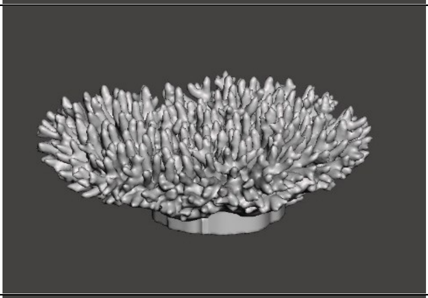 | Growth form with small closely woven branches with lots of secondary branching; often forming flat-topped clumps |
| **Digitate** | 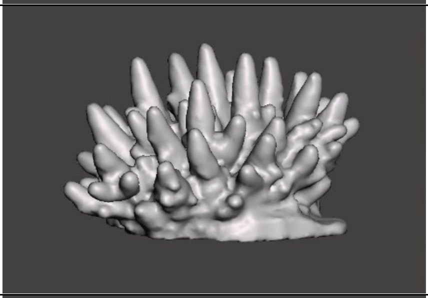 | Growth form with thick upright branches extending from a thick plate or encrusting base; without significant secondary branching |
| **Massive** | 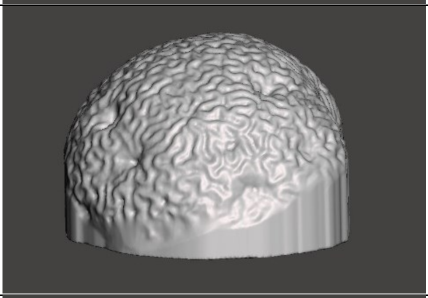 | Solid, boulder-like growth form, with a similar shape in all directions |
| **Tabular** | 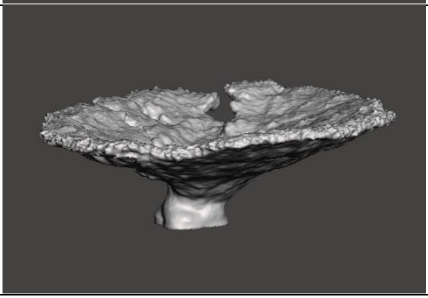 | Growth form with flat-topped platforms, often made up of short branchlets; typically, atop a singular central leg |

**SI Table 3** Metadata summary table of coral colony meshes extracted and used in the data analysis of this manuscript

| Growth form | Total number of meshes | Number of colonies (by number of years tracked) | | | | | Min planar area (cm²) | Max planar area (cm²) | Species | Number of species |
|---|---|---|---|---|---|---|---|---|---|---|
| | | 1 | 2 | 3 | 4 | 5 | | | | |
| Arborescent | 35 | 10 | 7 | 1 | 2 | 0 | 2.58 | 2067.30 | *Acropora affinis*<br>*Acropora donei*<br>*Acropora muricata*<br>*Acropora pacifica*<br>*Acropora vanderhorsti*<br>*Acropora yongei*<br>*Hydnophora rigida* | 7 |
| Branching closed | 41 | 0 | 3 | 4 | 2 | 3 | 8.29 | 1671.90 | *Pocillopora meandrina*<br>*Pocillopora verrucosa* | 2 |
| Corymbose | 63 | 4 | 11 | 7 | 4 | 0 | 1.03 | 1616.73 | *Acropora dissimilis*<br>*Acropora divaricata*<br>*Acropora erinae*<br>*Acropora kenti*<br>*Acropora nasuta*<br>*Acropora sarmentosa*<br>*Acropora* "shiny"<br>*(sensu Bridge)*<br>*Acropora spathulata* | 8 |
| Digitate | 50 | 0 | 6 | 3 | 6 | 1 | 5.45 | 323.01 | *Acropora* "fat" *digitifera*<br>*(sensu Wolstenholme)*<br>*Acropora gemmifera* | 2 |
| Massive | 99 | 14 | 8 | 8 | 5 | 5 | 4.24 | 1147.69 | *Dipsastraea matthaii*<br>*Dipsastraea pallida*<br>*Dipsastraea truncata*<br>*Favites complanata*<br>*Goniastrea retiformis*<br>*Hydnophora microconos*<br>*Leptoria phrygia*<br>*Lobophyllia recta*<br>*Paragoniastrea russelli*<br>*Platygyra daedalea*<br>*Porites* spp.<br>*Psammocora profundacella* | 12 |
| Tabular | 52 | 1 | 8 | 5 | 5 | 0 | 6.17 | 2565.28 | *Acropora bifurcata*<br>*Acropora hyacinthus*<br>*Acropora pectinata*<br>*Acropora tersa* | 4 |
| **Total** | **340** | 29 | 43 | 28 | 24 | 9 | 1.03 | 2565.28 | | **35** |

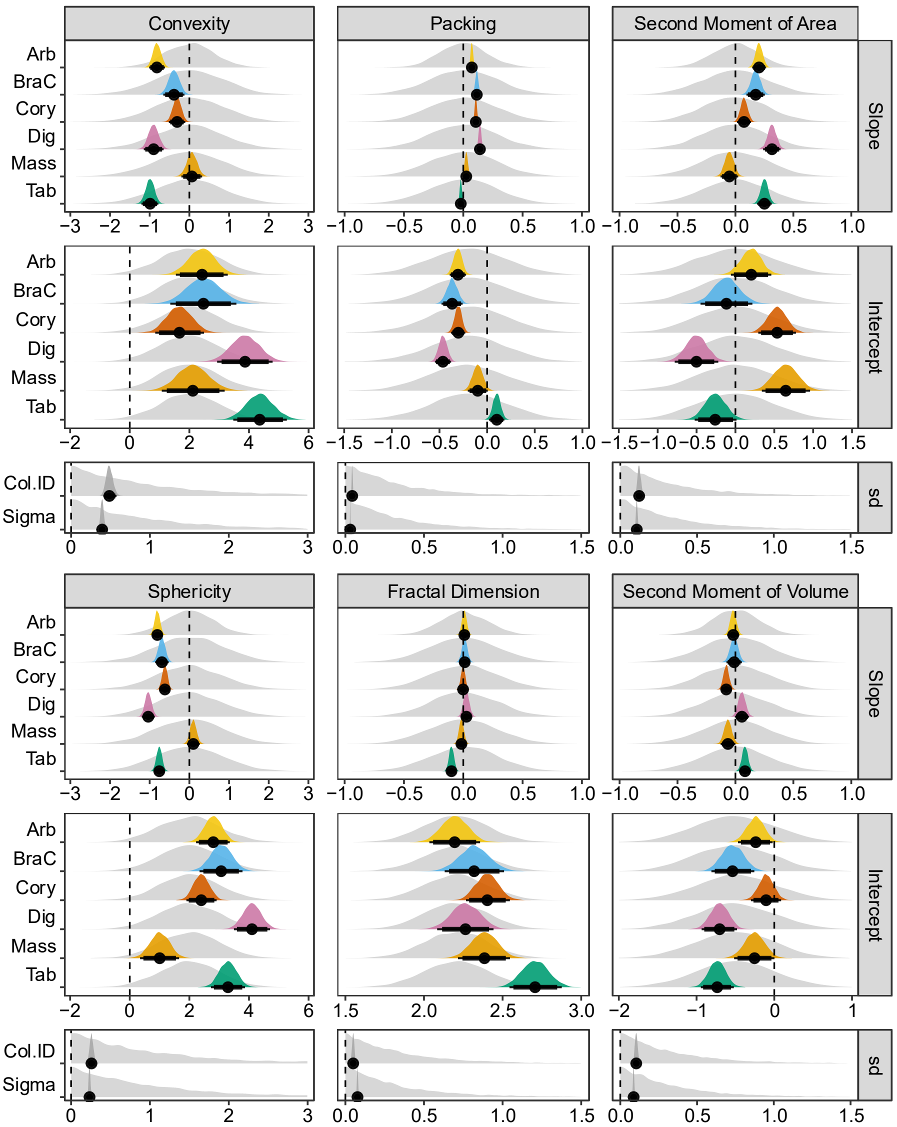


**SI Fig. 1** Prior predictive checks for slope, intercept, and standard deviation estimates for size-metric models. Light grey represents posterior distributions from the models run only on priors. Colored and dark grey distributions represent the half-eye plots of the posterior distributions run on the data, with the black bars depicting credible intervals around the median (thick = 90% credible interval, thin = 95% credible interval). Plots are gridded by model (horizontal) and estimates (vertical). Slope and intercept estimates are displayed by growth form (color) (Arb = arborescent, BraC = branching closed, Cory = corymbose, Dig = digitate, Mass = massive, Tab = tabular) and standard deviation estimates by fixed effects (dark grey) (Col.ID = Colony ID). A vertical dashed line (black) marks the 0

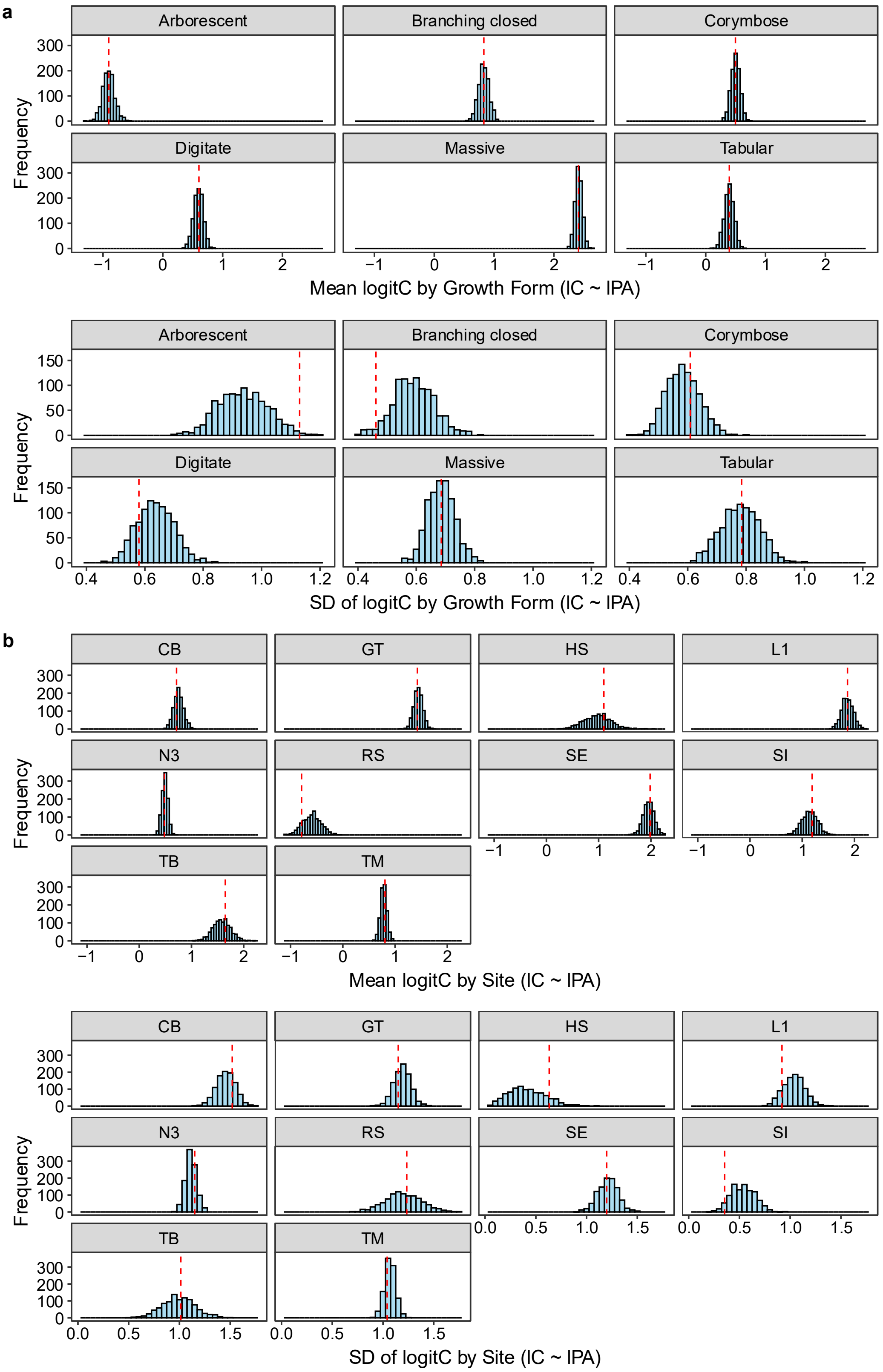


**SI Fig. 2** Posterior predictive checks for the mean and standard deviation (SD) of logit(convexity) values by growth form (a) and site (b) for the size-convexity model (lC ~ lPA). Histograms depict the distribution of test statistics mean (top) and standard deviation (bottom) computed for each of 1000 datasets simulated from the posterior predictive distribution. Vertical dashed lines (red) represent the test statistic computed from the raw data

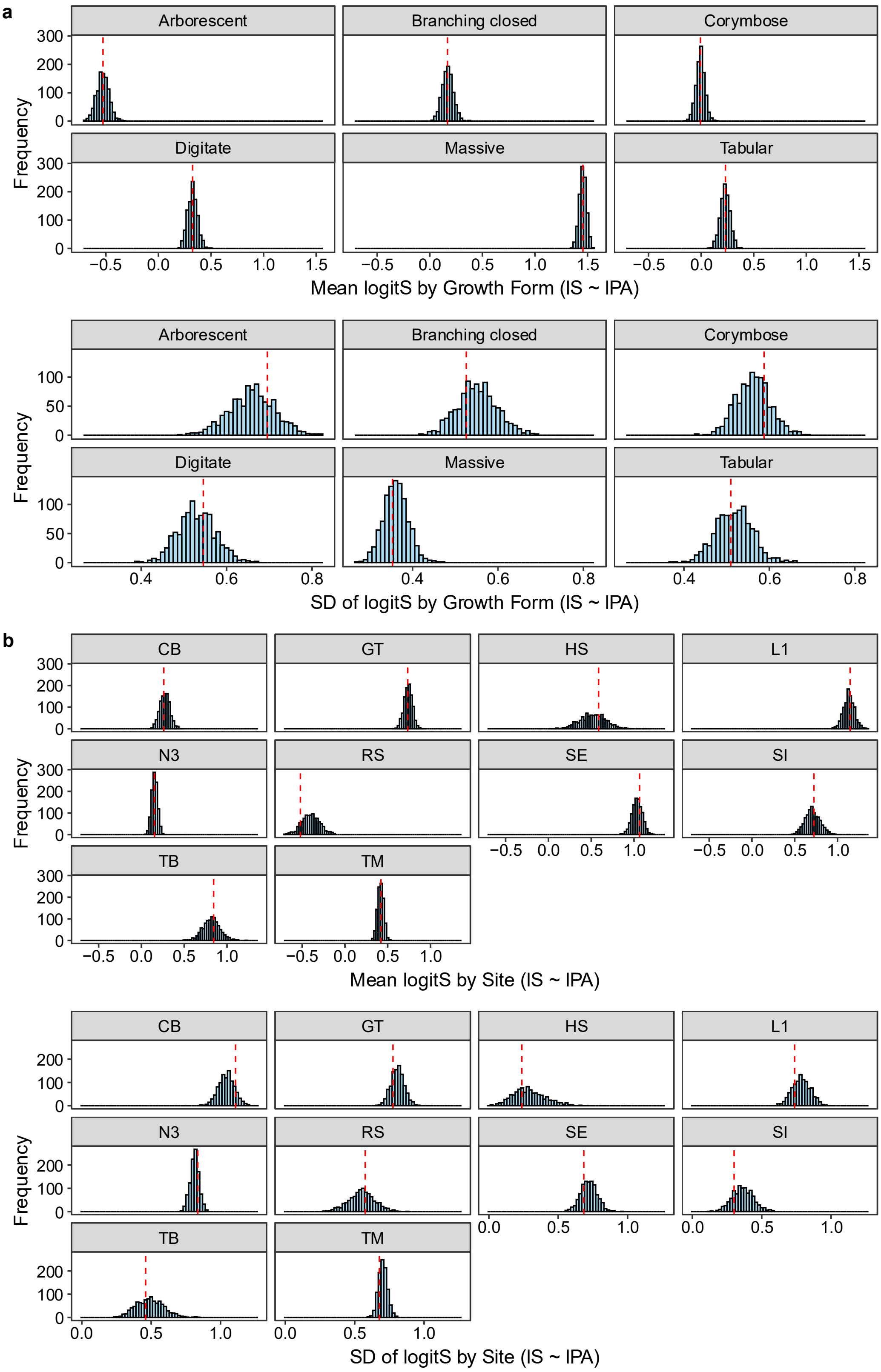


**SI Fig. 3** Posterior predictive checks for the mean and standard deviation (SD) of logit(sphericity) values by growth form (a) and site (b) for the size-sphericity model (lS ~ lPA). Histograms depict the distribution of test statistics mean (top) and standard deviation (bottom) computed for each of 1000 datasets simulated from the posterior predictive distribution. Vertical dashed lines (red) represent the test statistic computed from the raw data

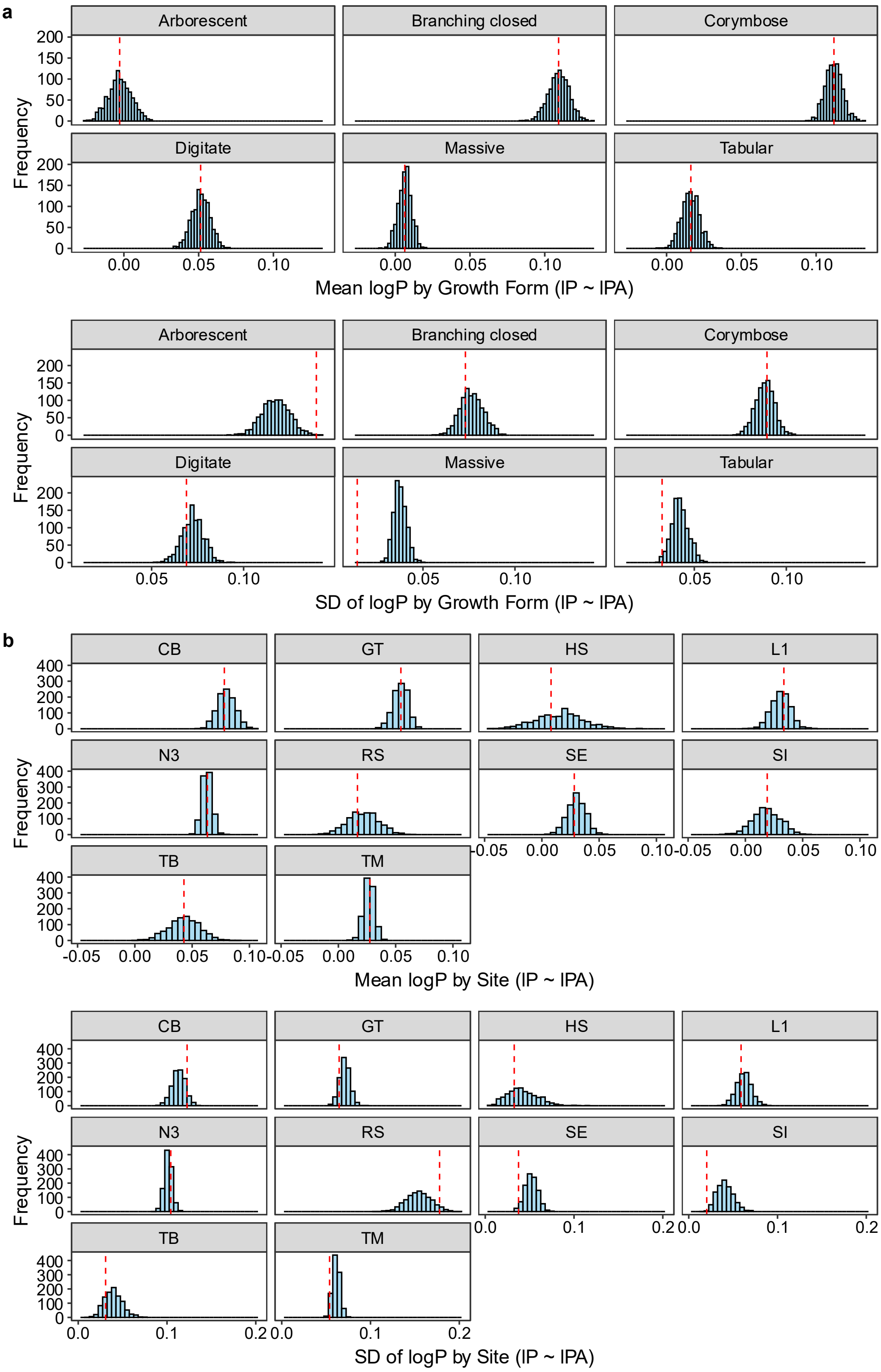


**SI Fig. 4** Posterior predictive checks for the mean and standard deviation (SD) of log(packing) values by growth form (a) and site (b) for the size-packing model (lP ~ lPA). Histograms depict the distribution of test statistics mean (top) and standard deviation (bottom) computed for each of 1000 datasets simulated from the posterior predictive distribution. Vertical dashed lines (red) represent the test statistic computed from the raw data

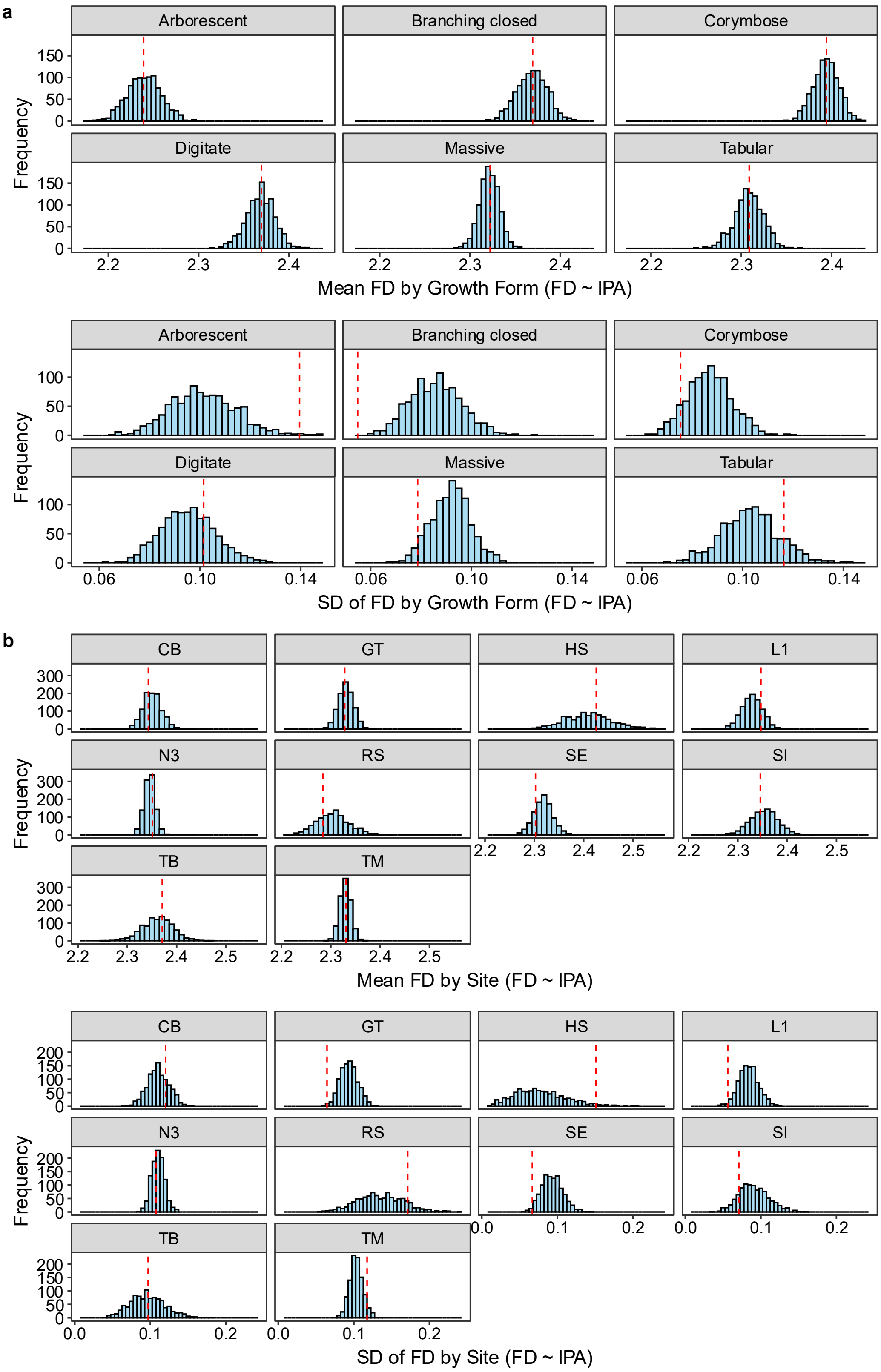


**SI Fig. 5** Posterior predictive checks for the mean and standard deviation (SD) of fractal dimension values by growth form (a) and site (b) for the size-fractal dimension model (FD ~ lPA). Histograms depict the distribution of test statistics mean (top) and standard deviation (bottom) computed for each of 1000 datasets simulated from the posterior predictive distribution. Vertical dashed lines (red) represent the test statistic computed from the raw data

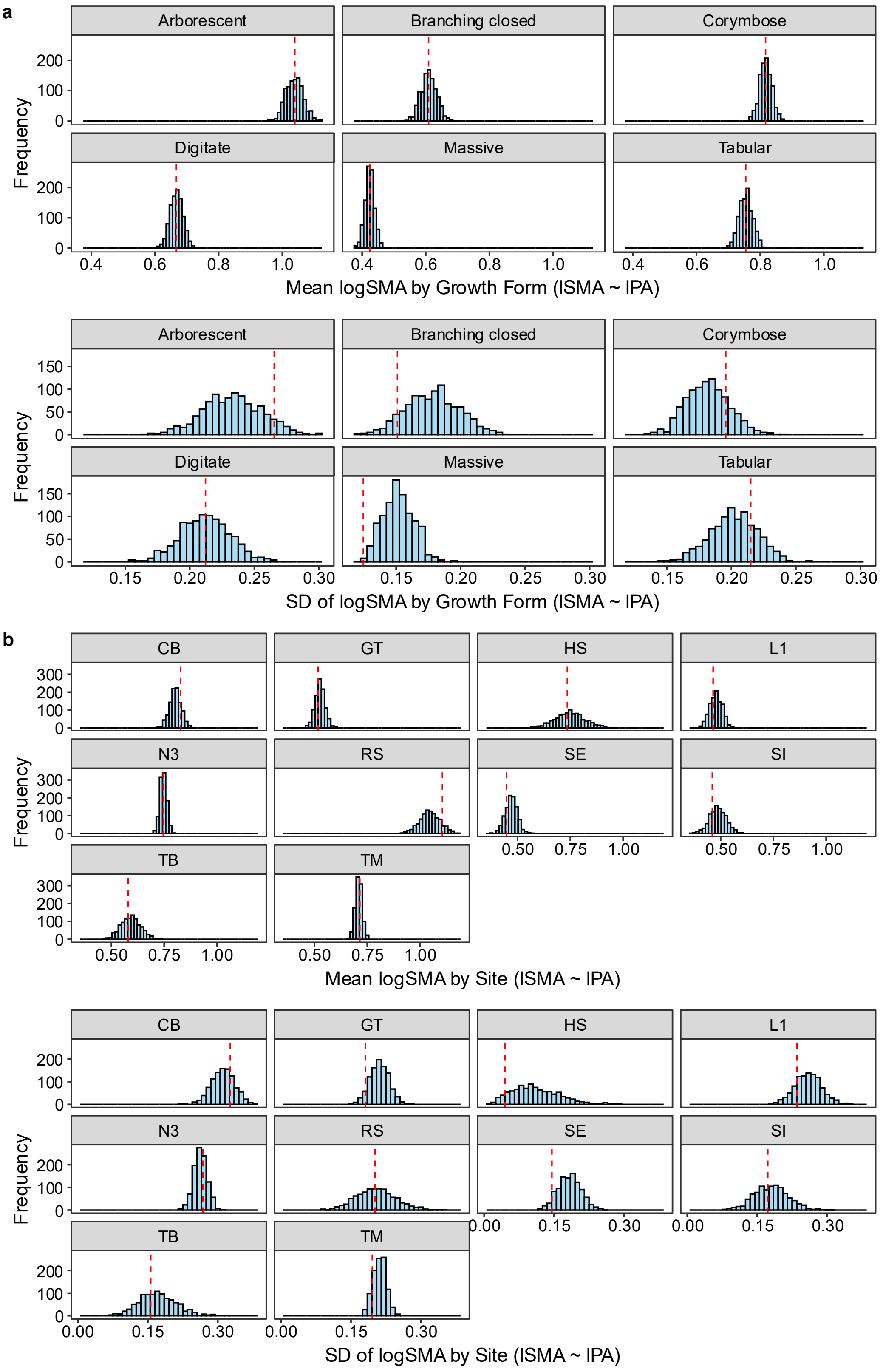


**SI Fig. 6** Posterior predictive checks for the mean and standard deviation (SD) of log(second moment of area) values by growth form (a) and site (b) for the size-second moment of area model (lSMA ~ lPA). Histograms depict the distribution of test statistics mean (top) and standard deviation (bottom) computed for each of 1000 datasets simulated from the posterior predictive distribution. Vertical dashed lines represent the test statistic computed from the raw data

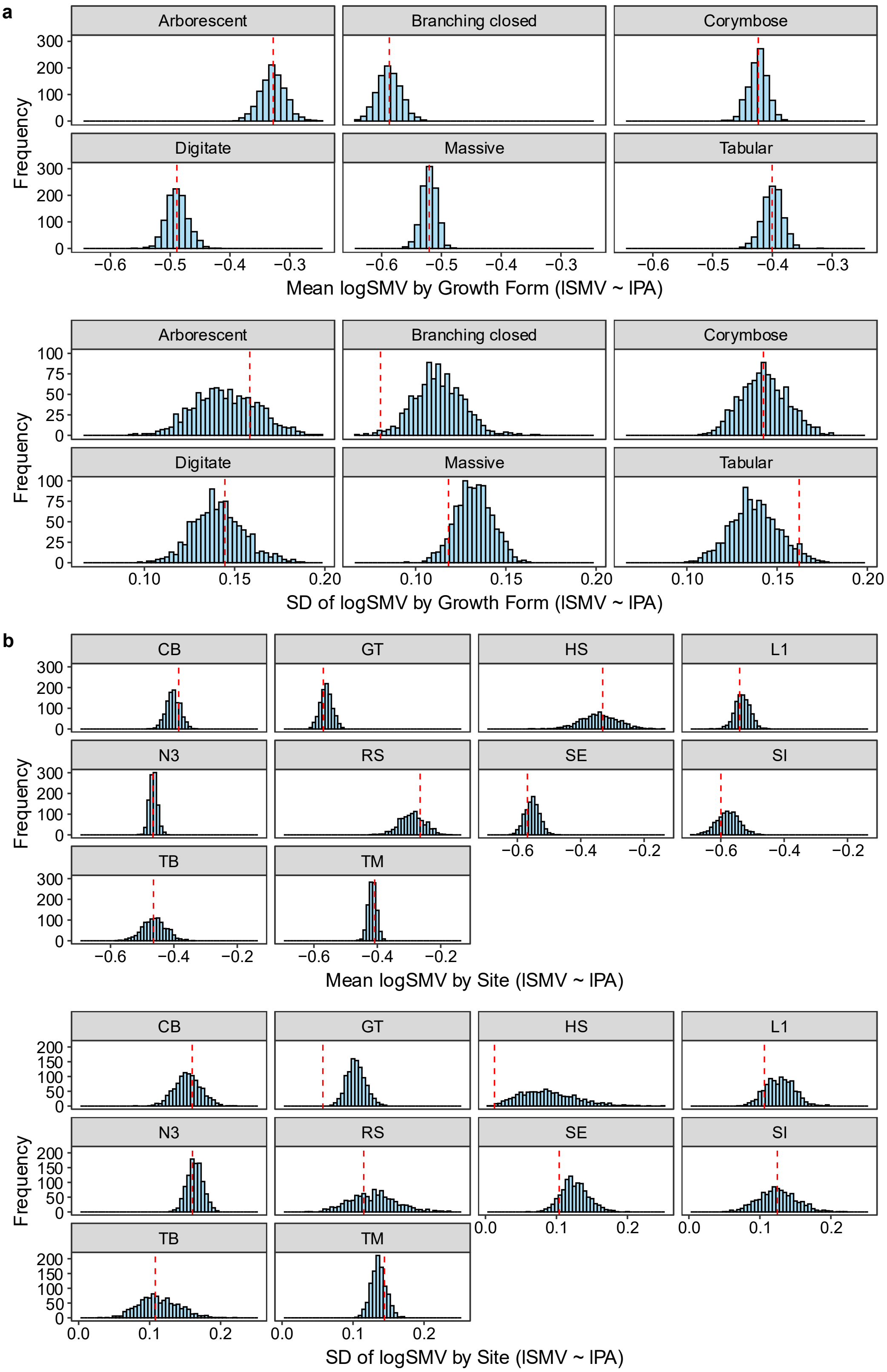


**SI Fig. 7** Posterior predictive checks for the mean and standard deviation (SD) of log(second moment of volume) values by growth form (a) and site (b) for the size-second moment of volume model (lSMV ~ lPA). Histograms depict the distribution of test statistics mean (top) and standard deviation (bottom) computed for each of 1000 datasets simulated from the posterior predictive distribution. Vertical dashed lines represent the test statistic computed from the raw data

**SI Table 4** Slope estimates (95% credible interval) by model across growth forms. Slope estimate intervals that do not cross zero are bolded

| Size-metric Model | Arborescent | Branching closed | Corymbose | Digitate | Massive | Tabular |
|---|---|---|---|---|---|---|
| Convexity | **-1.015 - -0.614** | **-0.656 - -0.124** | **-0.518 - -0.102** | **-1.149 - -0.649** | -0.201 - 0.323 | **-1.208 - -0.781** |
| Sphericity | **-0.939 - -0.673** | **-0.864 - -0.521** | **-0.745 - -0.492** | **-1.199 - -0.870** | -0.062 - 0.263 | **-0.898 - -0.620** |
| Packing | **0.052 - 0.092** | **0.088 - 0.141** | **0.087 - 0.125** | **0.117 - 0.163** | -0.001 - 0.051 | **-0.040 - -0.002** |
| Fractal Dimension | -0.029 - 0.048 | -0.034 - 0.057 | -0.038 - 0.033 | -0.022 - 0.075 | -0.056 - 0.025 | **-0.140 - -0.060** |
| Second Moment of Area | **0.144 - 0.265** | **0.095 - 0.254** | **0.014 - 0.135** | **0.239 - 0.390** | -0.129 - 0.024 | **0.187 - 0.314** |
| Second Moment of Volume | -0.070 - 0.033 | -0.077 - 0.056 | **-0.126 - -0.031** | -0.005 - 0.118 | **-0.128 - -0.000** | **0.032 - 0.133** |

**SI Table 5** Colony-level developmental trajectory magnitude and direction distributions (mean and 95%credible interval) by growth form

| Morphology | n | Magnitude | Direction (°N) |
|---|---|---|---|
| Arborescent | 60000 | 5.33 (4.60-6.07) | 120.4 (109.2-131.1) |
| Branching closed | 72000 | 5.49 (4.47-6.52) | 135.1 (122.1-147.3) |
| Corymbose | 132000 | 4.57 (3.75-5.39) | 149.2 (138.0-159.3) |
| Digitate | 96000 | 8.27 (7.41-9.19) | 122.8 (113.9-131.0) |
| Massive | 156000 | 1.42 (0.43-2.51) | 229.9 (184.6-289.2) |
| Tabular | 108000 | 6.16 (5.29-7.04) | 61.9 (53.6-70.7) |
